# Neural Interactome: Interactive Simulation of a Neuronal System

**DOI:** 10.1101/209155

**Authors:** Jimin Kim, William Leahy, Eli Shlizerman

## Abstract

Both connectivity and biophysical processes determine the functionality of neuronal networks. We, therefore, develop a real-time framework, called Neural Interactome^1^, to simultaneously visualize and interact with the structure and dynamics of such networks. Neural Interactome is a cross-platform framework, which combines graph visualization with the simulation of neural dynamics, or experimentally recorded multi neural time series, to allow application of stimuli to neurons to examine network responses. In addition, Neural Interactome supports structural changes, such as disconnection of neurons from the network (ablation feature), as typically done in experiments. Neural dynamics can be explored on a single neuron level (using a zoom feature), back in time (using a review feature) and recorded (using presets feature). We implement the framework using a model of the nervous system of Caenorhabditis elegans (C. elegans) nematode, a model organism for which full connectome and neural dynamics have been resolved. We show that Neural Interactome assists in studying neural response patterns associated with locomotion and other stimuli. In particular, we demonstrate how stimulation and ablation help in identifying neurons that shape particular dynamics. We examine scenarios that were experimentally studied, such as touch response circuit, and explore new scenarios that did not undergo elaborate experimental studies. The development of the Neural Interactome was guided by generic concepts to be applicable to neuronal networks with different neural connectivity and dynamics.

**Author Summary:** Emerging neuroimaging techniques and novel optical interfaces which record and control neural dynamics enable detailed computational connectivity and dynamics models for neurobiological systems. An open question stemming from these advances is how to validate, simulate and apply these models to predict network functionality. Supervised empirical exploration to identify functional stimulations is an elaborate process, and direct computational approach of sequential stimulation is also formidable since produces large amounts of data without clarity on how it can be used to steer toward meaningful functionalities. We therefore develop a platform to inspect network dynamics in real time while preserving structural connectivity properties, displaying the dynamics on a graph, with possibilities to identify functional sub circuits and review the simulated dynamics. The platform allows for real time interactions with the network such as variation of stimuli and performing connectivity changes as neural ablation. We apply the platform to Caenorhabditis elegans nematode nervous system model. We revisit experimentally known scenarios of stimulations and show how our platform helps to detect associated neural dynamic patterns within seconds through few interactions. In addition, we show how the platform could provide novel hypotheses for scenarios that were not yet explored empirically. By implementing the platform with flexibility for changes in connectivity and dynamic models, this work sets forth a generic methodology applicable to various neurobiological systems.

## Introduction

Modeling neuronal systems involves incorporating two modeling layers. The first fundamental layer is of neuronal connections static map (connectome). The layer on top of it is of biophysical processes of neural responses and interactions. In the recent years there has been significant progress in resolving and modeling both layers. Connectomes of several organisms and systems, such as the nematode *Caenorhabditis elegans (C. elegans)*, the *Drosophila* medulla, the mouse retina, mouse primary visual cortex, and others have been fully or partially mapped on various scales: from macro to single neuron level [1]–[6]. Also, decades of research in describing and modeling biophysical processes have provided both experimental and computational foundations for modeling single neuron dynamics as well as synaptic and electric processes between neurons [7]–[15]. Due to these advances, models incorporating both layers become more detailed and realizable for several neuronal systems. These models are called ***Dynomes*** as they correspond to dynamical system acting on top of the static connectome [16].

Being closer to the realistic neuronal system, *dynome* studies have more potential to reveal neural pathways and functionalities of the network [17], [18]. However, they also introduce challenges in finding appropriate methods for efficient studies of network capabilities [19]. Brute force approaches will typically produce formidable amounts of data, where extraction or characterization of relevant neural patterns can be cumbersome and time consuming. In particular, investigation of the *dynome* can be greatly enhanced by visualization that allows users to interact with network’s static and dynamic layers and to observe neural activity at the same time. In such a framework, the necessary components are (i) ability to apply or modify stimuli to the network as in experiments; (ii) being able to observe the neural dynamics on various time and population scales, and (iii) allow for network structural changes. Furthermore, the framework is expected to perform seamless integration for such functions and include review capabilities for exploration of the system and dynamics in depth. In this work, we thereby develop the Neural Interactome, which is a generalized visualization framework incorporating such capabilities. The framework employs a graph visualization layout to represent the static connectome. On top of the layout, it incorporates dynamic visual components to represent real-time neural responses. These components are implemented via synchronization method between the backend neural integrator of the *dynome* and the graph layout of the interactive interface. The backend neural integrator is connected to neurons stimuli panel, and permits setting external stimuli and changing the structure of the graph on demand. The framework also includes review and preset modes that allow for further exploration of simulated dynamics. In this paper, we focus on applying the framework to the nervous system of *Caenorhabditis elegans (C. elegans)* nematode.

*C. elegans* is a model organism for *dynome studies*, as it consists of 302 neurons with three types (sensory, inter, motor). Such a system is thus relatively small to be fully reconstructed and analyzed. Indeed, the near complete connectome of the nervous system has been resolved using serial section electron microscopy [3], [4], [20]. The connectome data includes enumeration of neural connections for the complete somatic nervous system (279 neurons) of ***synaptic*** type, where ***GABAergic*** neurons make inhibitory synapses, and ***glutamergic*** and ***cholinergic*** neurons form excitatory synapses. The connectome also enumerates ***gap*** junctions (electrical connections) for each pair of neurons. The connectome data is robust, since C. elegans neurons are recognizable and consistent throughout individual worms [3]. Furthermore, *C. elegans’* synaptic and gap connections are common across animals with more than 75% reproducibility [3], [21]-[23]. In addition to the anatomical structure of the nervous system, biophysical in-situ recordings of membrane voltage response to input current injected into each individual neuron in the network have been performed [11]. These revealed that *C. elegans* neurons are of non spiking type with graded potential membrane voltage profile [24]. Following these studies, a set of mathematical models describing neural membrane voltage and interaction between the neurons were developed [10], [24].

The availability of near-complete connectome data along with experimental quantification of responses and interactions provided a computational basis for reconstructing both static and dynamic layers of *C. elegans* neuronal network. Combination of these two layers was recently developed and constitutes the dynome: dynamical model for *C. elegans’* nervous system [10]. When applied with prescribed input stimuli, *C. elegans dynome* was capable of producing various forms of characteristic dynamics such as static, oscillatory, non oscillatory and transient voltage patterns. These dynamics indicated that *C. elegans dynome* is a valuable model for the worm’s nervous system, and patterns observed are suggested to be consistent with the experimentally observed ones. In particular, stimulation of sensory PLM neurons with constant current resulted in a two-mode dominant oscillatory behavior in forward locomotion motor neurons [10]. The model is expected to include a variety of other additional patterns, however, their full validation is formidable to perform, as it requires many simulations with various stimuli amplitudes and combinations. For instance, in the context of touch response, it would be valuable to examine stimulation of ALM and AVM sensory neurons, which in experiments was identified as associated with anterior touch response and expressed as backward crawling [25], [38], [39]. Furthermore, transitions from one type of dynamics to another (e.g. from oscillatory to non-oscillatory) are also expected to exist when input stimuli shift from one value to another. It is thereby introduction of a framework that eases these studies can assist in the validation goal. In addition, it can be used to ask a set of innovative questions that can provide valuable insights about *C. elegans* network.

**Fig 1.**
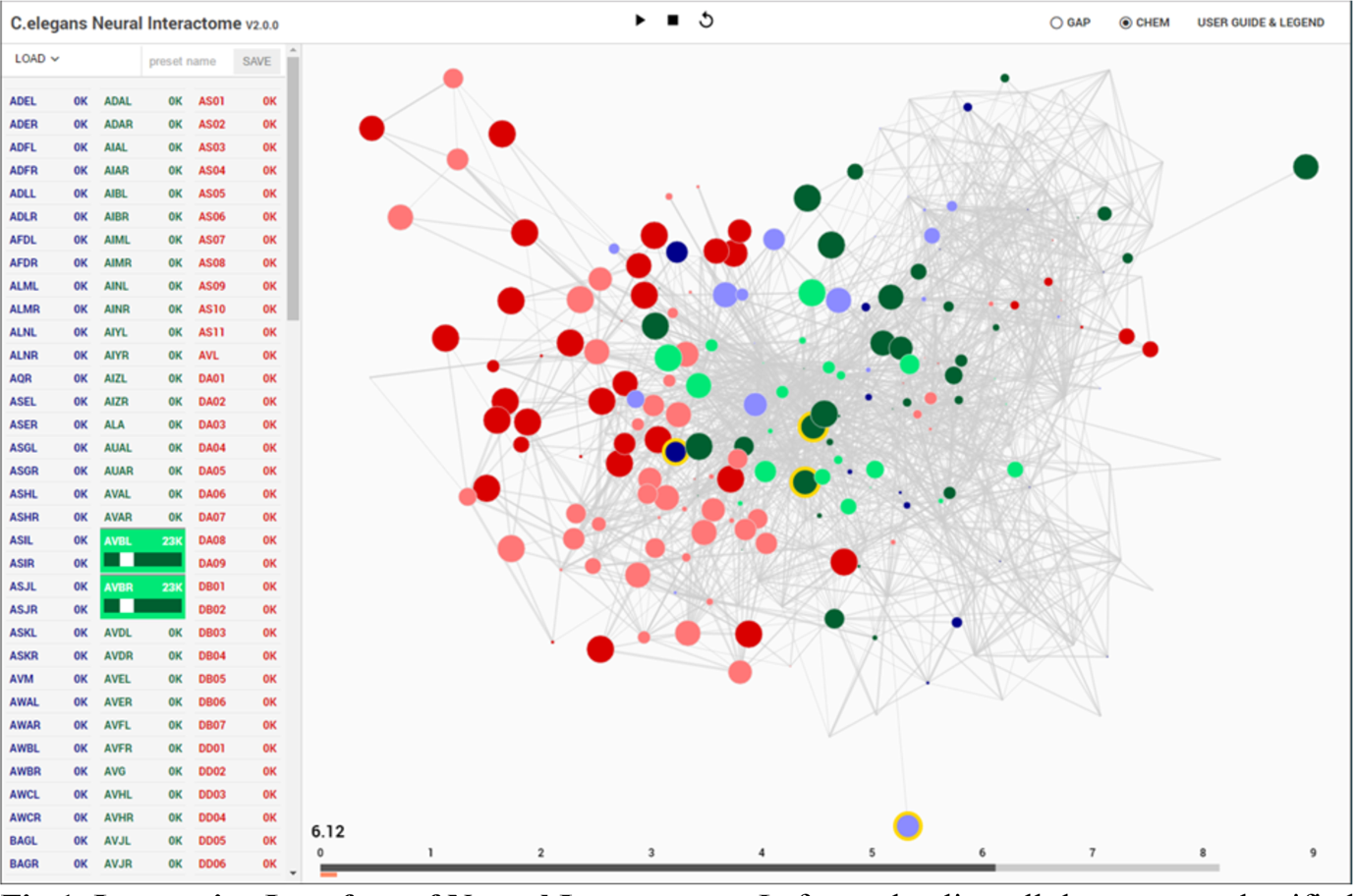
Interactive Interface of Neural Interactome. Left panel enlists all the neurons classified by type (sensory, inter and motor). Each neuron is a clickable button with a scroll option. Scrolling adjusts the magnitude of constant stimulus; click+shift ablates the neuron from the network. Right: Force-directed graph displays each neuron’s membrane voltage (node color denotes the sign and radius denotes the magnitude) and connections between neurons (edges between each pair of nodes). At the bottom of the graph, time bar keeps track of visualization current time (dark gray), and of computed time by the backend neural integration (light gray).

## Results

We first describe the main components of the Neural Interactome framework, and then continue to demonstrate its application to the nervous system of *C.elegans* worm for stimulation scenarios.

### Interactive Interface for Neuronal Network

Interactive interface for Neural Interactome consists of four main parts: (i) neural stimulation/ablation, (ii) visualization of *dynome* dynamics, (iii) control of simulation timescale, and (iv) review system.

#### Neural Stimulation & Ablation

Stimuli panel located on the left side of the screen controls neural stimulation (Fig. 1). It enlists all neurons in the network categorized into three group types (sensory, inter, motor). Each group type is given a characteristic color (sensory: blue, inter: green, motor: red). Each individual neuron on the panel is a clickable button with a scrollable bar, which allows setting amplitudes of constant stimulus, i.e., inject current to the neuron. The amplitude of the stimulus can be adjusted prior to running a simulation (as initial condition), or at any time during the simulation. When stimulus is being adjusted during the simulation, it effectively imitates ‘clamping’ of neurons in the network. In addition, to allow for testing various structural configurations for the network, the panel is designed to support neural ablation of neurons. By clicking on a neuron while holding the shift key, the neuron is grayed-out in the interface. Such operation disconnects the neuron from all of its respective connections (both receiving and outgoing) in both synaptic and gap type and thus effectively removes it from the network. The ablation can also be undone (reinsertion) by repeating the operation of ‘shift’ key + clicking on the ablated neuron. Similar to neural stimulation, both ablation and reinsertion can be performed both prior and during network simulation.

#### Dynome Visualization

##### Connectivity Representation

Visualization of *dynome* dynamics is on the right side of the interactive interface, with the connectome of the neuronal network represented as a graph (Fig. 1). The nodes of the graph represent neurons, whereas the edges represent connections (either gap or synaptic) between each pair of neurons. The top panel of Fig. 2 shows *C. elegans’* synaptic connectome (left) as well as its gap connectome (right), where each node represents an individual neuron and colored according to its group type. Initially, prior to displaying the *dynome* dynamics, the radii of the nodes are set according to in/out synaptic degree of the respective neuron (i.e. the amount of synaptic connections of a neuron). Such visualization emphasizes neurons with higher degree (hub neurons) by displaying them with larger radius and de-emphasizes neurons with lower degree with smaller radius. The width of the edge between a pair of neurons is set according to maximum synaptic weight, such that for a pair of neurons A and B, 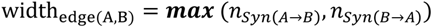, where 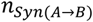 is the number of synapses from neuron A to B.

In addition, we use force-directed graph algorithm to arrange the nodes and edges in optimal positions [26]. The algorithm visualizes graphs by assigning forces to nodes and edges based on their relative positions and routings. For edges, spring-like attractive forces based on Hooke’s law are used to attract pairs of endpoints towards each other. For the nodes, repulsive forces, e.g. Coulomb’s law forces, are used to separate all pairs of nodes. Once forces are assigned, the algorithm minimizes the total energy potential of the system (i.e. equilibrium states for the system of forces) and displays optimal nodes and edges configuration. In this representation, the edges tend to have uniform lengths (due to spring forces) and the nodes that aren’t connected by an edge tend to be drawn further apart (due to repulsive forces) [27]. We found such graph visualization advantageous for neuronal networks as it: (i) keeps approximately equal lengths for all neuron’s connections, and (ii) arranges the nodes such that neurons that make connections with a particular neuron are found within its proximity. We also found that due to these properties, the configuration depicts the network in an intuitive way, by grouping the same type of neurons together in the network (e.g. many of the motor neurons are clustered together on the left of the graph) and places the neurons with high synaptic degrees in the middle. To keep the same frame of reference, force-directed representation is pre-computed before the simulation such that the positions of the nodes remain constant at all times.

**Fig 2.**
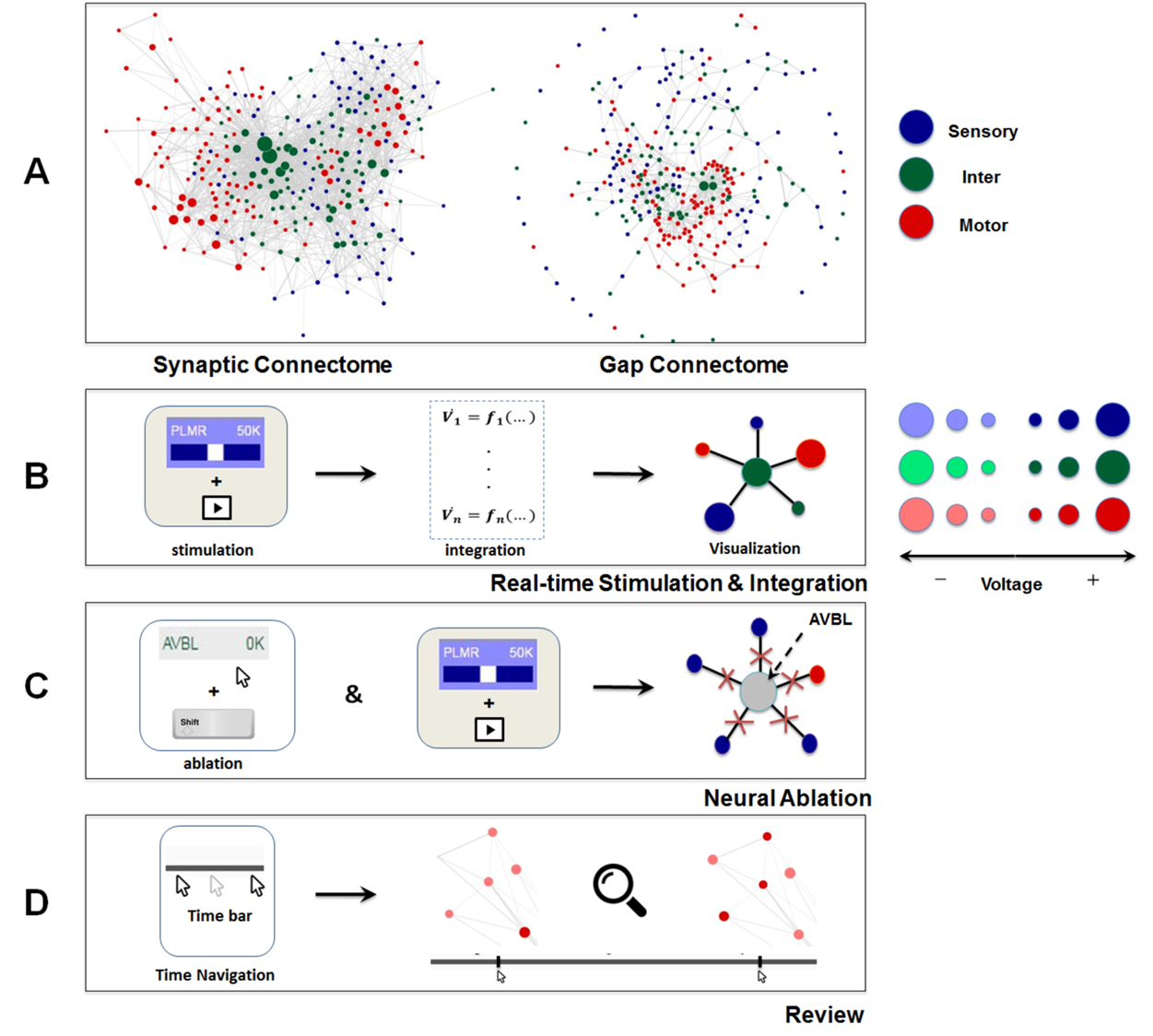
Dynome visualization and main functionalities of Neural Interactome. **A:** Force-directed graph visualization of *C.elegans* worm’s synaptic (chemical, left) and gap (electrical, right) connectomes. Each node represents individual neuron colored according to its group type and edges represent connections (either synaptic or gap). **B:** Schematics of Neural Interactome’s real-time stimulation and integration component; when user stimulates a neuron on the interface, the backend integrator computes membrane voltages in a response to the stimuli, which are then visualized on the graph in respect to their signs and magnitudes. **C:** Neural ablation is performed by clicking on the neuron, while holding the shift key. Ablation disconnects all the connections of the neuron (both gap and synaptic). **D:** The review system implements a clickable time bar to allow navigation to any previously computed time point in the simulated dynamics. This is further enhanced with dynamics zoom-in/zoom-out feature designed for in-depth analysis of connectivity structure and local dynamics of sub-circuits.

##### Neural Activity Visualization

Neural activity is represented as additional layer on top of the static connectivity graph. We find that optimal approach to visualize the two layers is to alter graph components (nodes and edges) according to neural activity. This creates a ‘breathing graph’ which represents network activity and structure in real-time. In particular we propose dynamically changing radii and colors of the nodes to depict neural activity. Changes are typically noticeable when the visualized variable representing the activity is continuous and scaled. In addition, it is beneficial that the visualized variable will have a biophysical meaning. Notable candidates for such variables are SR calcium activation dynamics or instantaneous firing rates (*peri stimulus time histograms* PSTH) [28], [29], [30]. SR calcium activation is a scalable continuous process representing transformation of membrane voltage dynamics (spiking, bursts separated by near-silent interburst periods, and graded voltage potentials) to an activation variable. It serves as a vital biophysical signal associated with activation of muscle activity [31]. In addition, several recording techniques quantifying neural dynamics are capable to measure and monitor SR calcium activity and can be directly compared with the visualization. The PSTH variable is computed from spiking dynamics and represents a spike count over a sliding window in time [32]. Such a measure is applied to both measurements of spiking membrane voltage or a computational model that produces spike trains. PSTH is a continuous and scaled measure widely used for identification, classification and recognition of response patterns associated with stimuli [33], [34].

To visualize these activity variables we propose to alter node radius and color. When the variable is signed (as in SR calcium activation) we use the radius to represent variable’s amplitude and assign a color map to represent its sign. When the variable is unsigned, as in the case of PSTH, only one node component (either color or radius) is needed to represent its amplitude, and the other component can be utilized for visualization of additional information such as spike times. For example, when the radius is used to depict the PSTH amplitude, color flickering can be used to display the occurrence of spikes.

For *C. elegans* network we transform membrane voltage to SR calcium like activation variable to represent neural dynamics. In particular, membrane voltages, computed by backend neural integrator, described further in ‘*Backend Neural Integration*, are translated to the following metric of radius size according to:

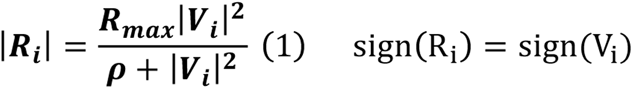

where *R_max_* is the maximum radii of the nodes and ρ is the slope factor. The sign of R_i_ is determined by the sign of the voltage V_i_. Such scaling of membrane voltages allows discerning active neurons at each given time without having to visualize the raw voltages. While in *C. elegans* membrane voltages are graded potentials, similar scaling accommodates other diverse types of neural activity, e.g. bursts, oscillations, etc [35].

Observing the colors and radii scaling over time allows to visually capture the unique patterns of dynamics on a population level, specifically oscillations, sudden bursts, settling down of dynamics. For example, in oscillatory dynamics of a population, colors will distinguish representatives of particular groups that are active and dynamically change their tones to display the fluctuation between positive and negative voltages. Indeed, for *C.elegans* network we show how we can identify oscillatory sub circuits of motor neurons, which fluctuate from negative to positive values over the period of 2 sec, upon stimulation of PLM touch sensitive sensory neurons.

##### Simulation Timescale

We implement the simulation timescale to be typically slower than the actual time in order to: (i) balance computations performed by the backend, and (ii) allow users to capture details of the dynamics, as visualization in actual timescale tends to happen too quickly. We also design the timescales of the stimulations to be dynamic, such that during stimuli transition or neural ablation, running time temporarily slows down to capture the dynamics that occur during the transition.

On the bottom of the interface we locate the time bar, which serves as the interface for users to observe the timescales of the visualization (Fig. 1). It consists of two bars; the dark gray bar shows the current time in visualization, while the light gray bar shows the computed time by the backend. We have developed the time bar to be similar to a streaming bar, which is widely implemented in popular video-hosting websites such as YouTube, and provides the interface to our *review system*, as we describe next.

#### Review System

The *review system* allows for isolating various time and population scales for further analysis (Fig. 2, Panel D). Using the time bar, we add the ability to navigate back to any previously computed time by clicking on a desired time point within the time bar (analogous to navigating back and forth while playing a video). In such a case the network along with the dark gray bar are reset to the state at the selected time point. Time navigation can be done either during simulation or when simulation is paused. In the former, the simulation will continue onward from the selected time point while for the latter, it will display the paused dynamics at that time point. In addition, we assign left and right arrow keys on the keyboard for the control of visualization speed (Fast FWD and Fast BWD). When activated during simulation or paused, the left and right arrow keys increase visualization speed while browsing through the dynamics in both directions.

Additional component of the *review system* for exploration of population scale is the dynamic zoom-in/out feature. The feature implements focusing into sub circuits within the network at any time during the simulation. It is implemented by keeping the nodes of same size and uniformly scaling the lengths of the edges. Effectively, such a method is optimal for observing a small group of neurons, as it increases the spacing between nodes and displays local sub circuit connectivity structure and dynamics (Fig. 2, Panel D). Hovering with a mouse over a neuron will also highlight the connections it makes to neighbor neurons, and display labels categorized in different group type colors.

We also incorporate presets feature as part of the *review system*, which allows users to save configurations of neurons stimuli panel. This feature is particularly useful when one wishes to save the stimuli and ablation setting, which led to interesting dynamics. Configuration can be saved by entering the desired name for the preset on top of the neurons stimuli panel (Fig. 1).

### Backend Neural Integration

Backend neural integration computes membrane voltage values for the whole network for a time interval [t, t + Δt] and transmits these values to the interactive interface for visualization. To compute the voltages, the integrator is solving a system of nonlinear ordinary differential equations that models the biophysical processes and interactions between neurons. For *C.elegans*, following equations are being integrated (see Kunert et al, 2014 for more details):

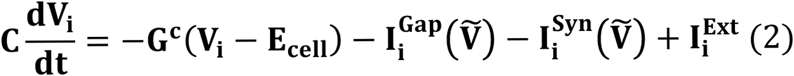

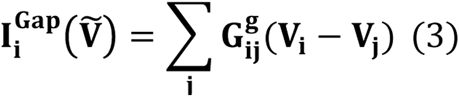

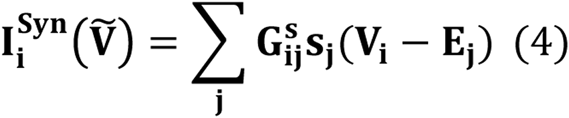

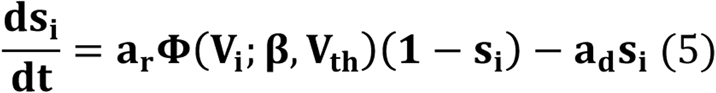

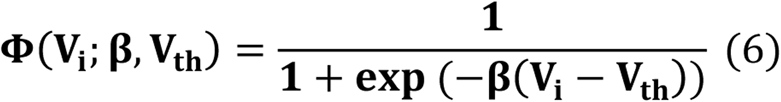

Where **C** is the cell membrane capacitance, *G^c^* is the cell membrane conductance, *E_cell_* is the leakage potential, and 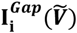, 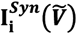, and 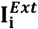 each correspond to input current contributed by gap junctions, synapses, and external input stimuli. Note that synaptic current is proportional to the reversal potential **E_j_**. 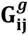 and 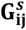 each correspond to total conductivity of the gap junctions between i and j and maximum total conductivity of synapses from i to j, where 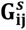 is modulated by synaptic activity variable **S_i_**. The synaptic activity variable is governed by equation (5), where **a_r_** and **a_d_** correspond to the synaptic activity rise and decay time, and **Φ** is the sigmoid function with width **β**.

**Fig 3.**
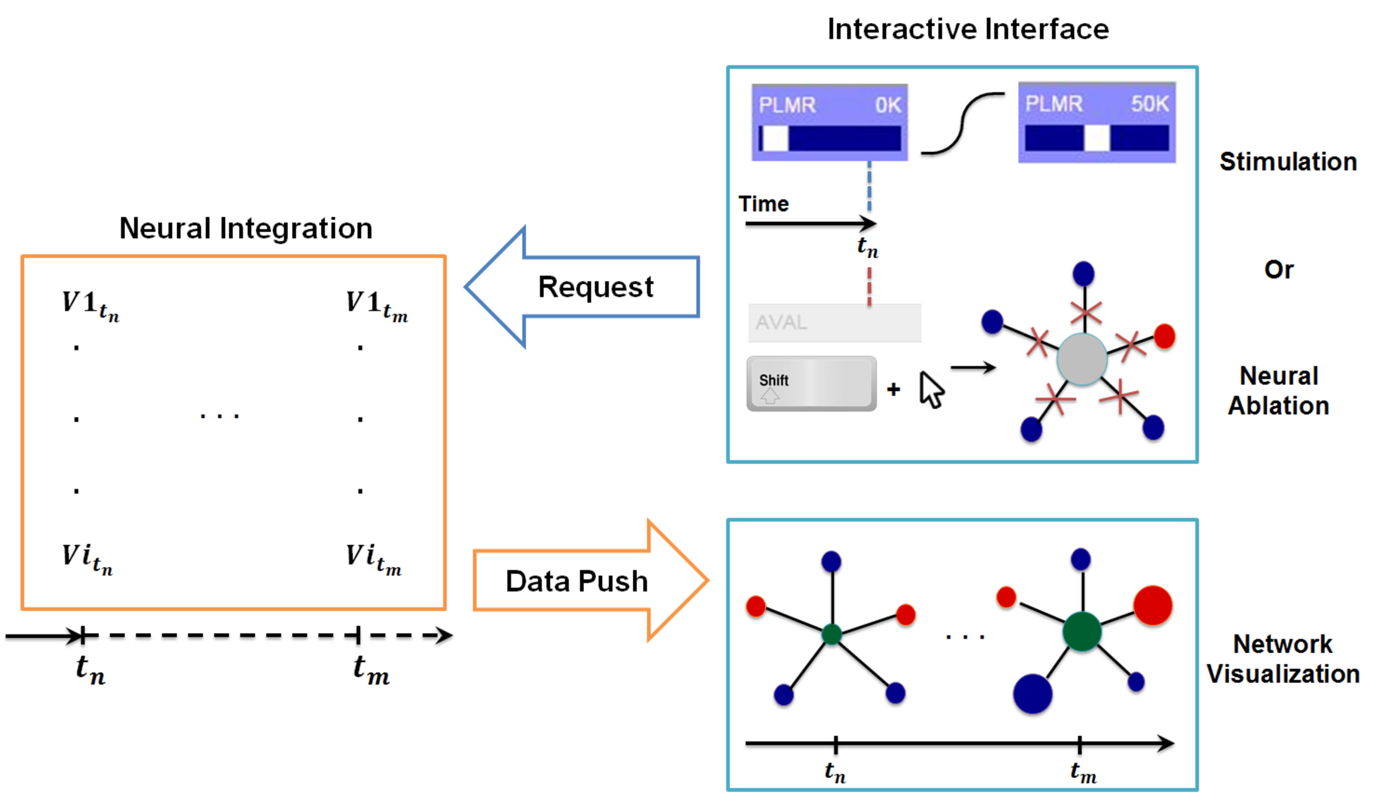
Synchronization between interactive interface and backend neural integration. The backend computes membrane voltage values for future time interval requested by the interface, and transmits them to the graph for be visualization. User driven change in the interactive interface, i.e. stimulation or neural ablation, invokes a process that passes the information to the backend where relevant parameters of integration are modified.

#### Synchronization of integration and visualization

For real-time interaction, we implement a synchronization procedure through a communication system between the interface and the backend. The protocol monitors the following quantities: **t_computed_**: Computed time at backend neural integration, **t_Visualization_**: Visualized time at interactive interface, **Δt**: Data stack, i.e. time interval to be computed, **t_buffer_**: Buffer size between t_computed_ and t_Visualization_, **τ**: Internal refractory period for checking t_computed_ — t_Visualization_.

The system is implemented to keep t_computed_ and t_Visualization_ ≅ t_buffer_ at all times such that backend neural integration is always responsive to real-time user interactions, but also accommodate computation of new solutions before the visualization fully catches up with the computation.

Based on these principles the communication protocol is as follows:

(i) The interface sends a command to the backend to compute solutions for the time interval of [t_computed_, t_computed_ + Δt] given the condition:

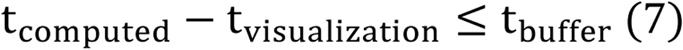
(ii) Once the command has been sent, the interface waits for a new block of solution of size Δt from the backend.
(iii) Once the block is received the interface resumes to poll whether condition (7) is satisfied. Polling is performed as follows: If the condition is met, the system applies (i). If not, the system goes through a refractory period of τ and then checks again for the condition (7).

In Fig. 3 we include a diagram of how the synchronization method allows for stimulation of neurons at any given time and simultaneous inspection of network response to such actions. When the user stimulates a specific neuron (e.g. PLMR in Fig. 3) or performs neural ablation, the interface sends a command to backend neural integration to modify necessary parameters. This is followed by an additional command from the interface to compute the solution for interval [t_stimulus_, t_stimulus_ + Δt]. The backend, upon receiving the first command, modifies the input stimuli parameters for stimulation or connectivity matrices for ablation. It then executes the second command by computing the voltage values for all neurons for a given time interval. The computed voltage values are then transmitted to interface for visualization. This cycle of command and data transmission is repeated indefinitely until the simulation is stopped.

#### Stimuli Transition

In addition to integration of the dynamical equations, the backend ensures that any modification of stimuli amplitude during the simulation is executed in realistic manner (i.e. no sudden jumps or drops in the stimulus). Ensuring such continuity produces more realistic shift of stimuli from one value to the other. Explicitly, we determine the magnitude of stimulus during the transition through combination of two the hyperbolic tangent functions:

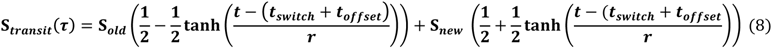

Where *t_switch_* is the time when the user injects or modify the input current, and **r, *t_offset_*** are the constant coefficients that determine the width and initial point of the transition respectively. Such construction makes sure that every transition takes place in a continuous manner and allows modifying the relative speed of transition by adjusting the value of **r**.

#### Neural Ablation

In addition, the backend implements neural ablation by instantaneous modification of connectivity matrices (both gap and synaptic). This step is followed by recalculation of the quantities in the network associated with a modified structure (e.g. the equilibrium states of the network Vth; See *Materials & Method* section for more detail). Effectively, when the user ablates a neuron in the interface, an array that keeps track of active neurons (1-present, 0-ablated) is being updated. The modified array is then sent to the backend, where for each ablated neuron, say neuron *i*, all elements of the connectivity matrices at row and column *i* (corresponding to in/out connections) are set to zero.

Reinsertion of neurons after they were ablated implements the ablation process in backward sequence of operations. Particularly, when the user reinserts the neuron, the interactive interface modifies the active neurons array, such that the corresponding neuron’s entry is changed from 0 to 1. The modified array is then transmitted to the backend, where it will restore the corresponding row and column of the connectivity matrices to default values.

### Scenarios: Locomotion & Nictation Stimuli

We proceed to demonstrate how application of Neural Interactome to *C.elegans* nervous system can assist in the study of neural dynamics. In particular, we target a sub circuit associated with a touch response, which stimulation is known to be associated with forward and backward locomotion. We also explore a neural dynamics pattern induced by the excitation of sub group of sensory neurons, which recently were discovered to be associated with nictation behavior.

#### Posterior touch response stimulation scenario

We stimulate PLM sensory neurons and AVB interneurons with constant stimuli to examine neural patterns associated with forward crawling motion as a result of posterior touch response. PLM sensory neurons (PLML/PLMR) in *C. elegans* nervous system are known as posterior mechanoreceptors. When stimulated by tail touch, PLM neurons excite motor neurons associated with forward crawling motion [25]. AVB interneurons (AVBL/AVBR) on the other hand, are part of the locomotory sub circuit and are also known as driver cells for forward movement of the worm. For the magnitudes of input currents, we adjust them by scrolling stimuli bars in the interface. Specifically, we set 1.4 * 10^4^ for PLM neurons, and 2.3 * 10^4^ for AVB interneurons, which result in profound oscillations.

As expected from experimental results and prior work, we observe oscillations in some populations of neurons. We therefore study their periodic cycle. In top panel of Fig. 4. Panel A, we show two snapshots of network dynamics taken at discrete percentages into the periodic cycle. We observe that the network graph responds with strong oscillation in about ~40% of the neurons with mostly motor neurons (marked in red) being specifically active.

We identify more detailed properties of the dynamics by inspecting the dynamic graph in review mode (Fig. 4. Panel B). The interface allows us to identify most responsive neurons and classify them into different types. In motor neurons most active neurons (by maximum voltage amplitude above the threshold) appear to be Ventricular and Dorsal type B (VB, DB) neurons alongside with Ventricular and Dorsal type D (VD, DD). Both of these types have identical oscillatory period of ~2 seconds, however, their dynamics are out of phase to one another. These observations re-confirm and clarify the oscillatory property of the dynamics reported in the literature and the most active neurons involved in such stimulation [25], [36], [37].

Most responsive interneurons turn out to be AVB, LUA, DVA, PVR and PVC. Indeed AVB, DVA and PVC were experimentally shown to act as modulators for forward locomotion [11], [25], [38], [39]. Notably, Neural Interactome also identifies relatively strong responses in LUA and PVR neurons. While these neurons have structural connections to PLM (LUA neurons are suggested to connect between PLM touch receptors, and PVR have gap junctions to PLM), their direct relation to forward locomotion was not affirmed (e.g. laser ablation of LUA did not lead to abnormalities of movement). Our analysis, however, suggests that these neurons are actively participating in the oscillations. These findings thus suggest that Neural Interactome can help find candidates of neurons correlated to particular dynamics, even for known sub circuits.

**Fig 4.**
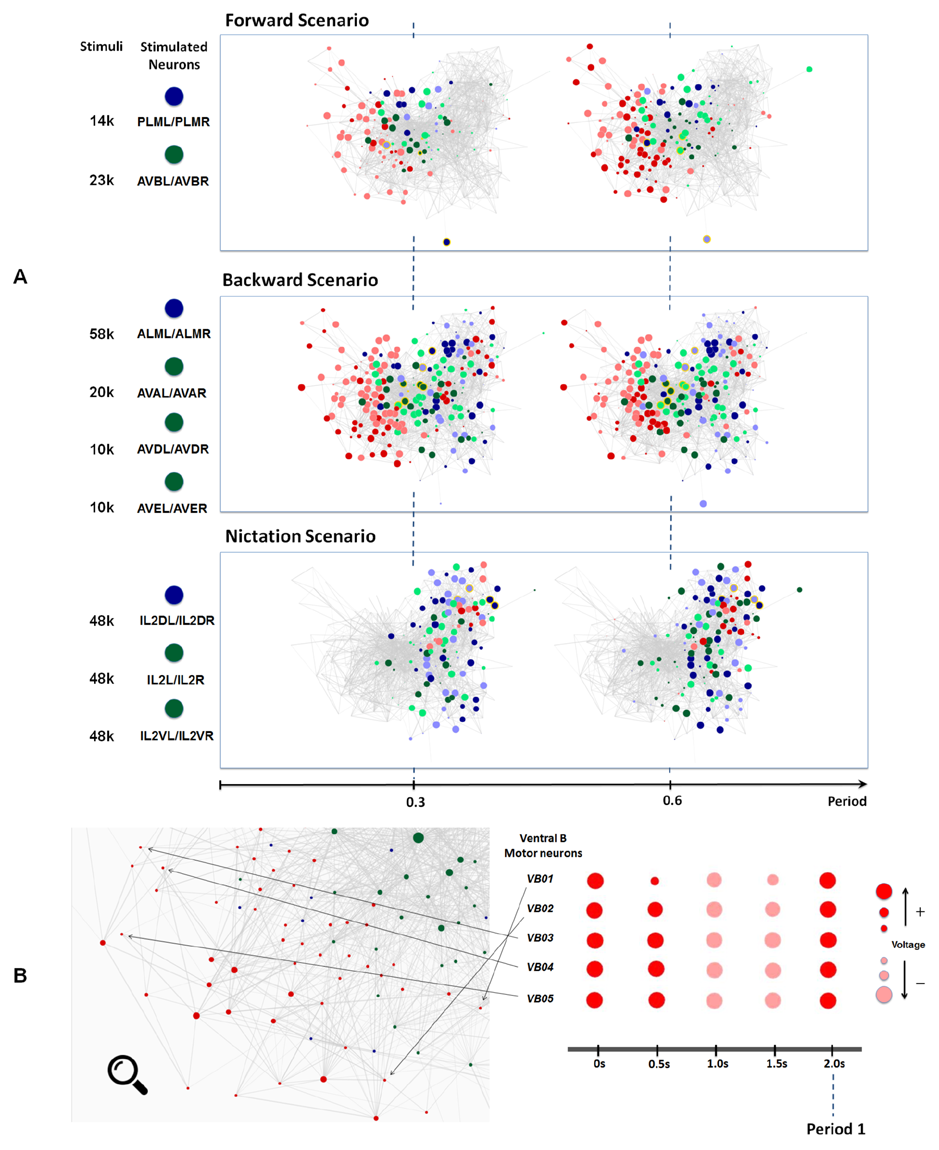
A. Snapshots of neuronal responses corresponding to forward, backward crawling and nictation scenarios visualized by Neural Interactome. Forward scenario shows the snapshots of neural dynamics in response to PLM and AVB neurons stimulation. Each snapshot is taken at 30% and 60% into the average period of motor neurons oscillatory dynamics. Backward scenario displays the snapshots of neural dynamics as a result of stimulation of ALM/AVA/AVD/AVE neurons, which are related to backward locomotion. Finally, Nictation scenario displays the snapshots of dynamics upon stimulation of IL2 neurons, which are known to induce a dispersal behavior. All scenarios are sampled at same percentages into average period of motor neurons oscillatory dynamics. **B. Identification of unique oscillatory dynamics during forward scenario using review mode**. Visualization of motor-neurons sub circuit using review system zoom-in function (left). Snapshots of Ventricular B motor neurons (VB01 ~ VB05) during forward scenario sampled five times with equal interval during 2 sec periodic cycle (right).

##### Anterior touch response stimulation scenario

For examination of neural patterns associated with anterior touch response, often triggering backward locomotion, we stimulate ALM sensory neurons along with AVA/AVD/AVE interneurons. ALM neurons sense touch to the anterior body region (i.e. frontal body) and induce motor neurons dynamics associated with backward locomotion [25]. Aiding this process are the AVA, AVD and AVE interneurons which act as modulators for the motion. We therefore stimulate these neurons with input currents that lead to profound dynamics, in particular: ALM = 5.8 * 10^4^, AVA = 2 * 10^4^, and AVD/AVE = 1 * 10^4^.

The snapshots of network dynamics while stimulating these neurons are presented in the middle panel of Fig. 4. Panel A and labeled as Backward Scenario. Notably, comparing forward vs. backward neural responses, the dynamics for backward involve much larger number of neurons than that of forward motion (~90%). We find the most responsive motor neurons to be Ventricular Dorsal type A (VA, DA), Ventricular Dorsal type D (VD, DD) and AS (AS01 ~ AS10). The oscillation behavior for each of these groups is of different phase, but their periods appear to be uniform around ~3.5 seconds. The results are consistent with the experimental observations which reported the A-type motor-neurons as directing cells along with D type motor neurons coordinating the backward motion [25], [37].

Zooming into particular populations of motor neurons we observe that individual motor neurons exhibit more complex and irregular patterns than those of the forward locomotion. Unlike the oscillations observed in forward motion which are characterized by predominantly smooth sinusoidal form, here motor neurons appear to have oscillatory patterns with varied waveforms: some motor neurons repeat aggressive fluctuations between negative and positive voltage while some exhibit triangular type oscillations above their thresholds. These results suggest that the motor-neurons pattern associated with backward crawling have different structure than of the forward crawling.

We also observe more activity within the interneurons. Most prominent ones appear to be AVA, AVDR, AVE, PVR, DVA, ADA and SABV. Some of these neurons are indeed identified in the literature, AVA, AVD, AVE are characterized to act as modulators for backward motion and DVA is characterized to maintain activity [11], [25], [38]-[40]. However, we also find high activity in neurons such as PVR, ADA and SABV. While PVR neuron makes gap junction with ALM, its role in backward locomotion has not been yet clarified. For both ADA and SABV, while SABV are known to innervate head muscles, their functionality has not been fully specified yet [3]. As in the posterior touch response scenario, the discovery of these additional neurons participating in dynamics provides new insights regarding the neurosensory integration of anterior touch response behavior.

##### IL2 neurons stimulation scenario

It has been recently shown that IL2 neurons regulate behavior called *nictation*, in which a worm stands on its tail and waves its head. Such behavior is known to be observed within *dauer* larva (i.e. developmental stage nematode worms), to transport itself to different niche via hosts such as flies or birds under harsh condition [41]. For the non-*dauers*, targeted activation of IL2 neurons does not induce nictation, possibly because IL2 neurons undergo a significant structural change at the *dauer* stage. In this scenario, we stimulate IL2 neurons through Neural Interactome, to investigate motor neuron dynamics possibly linked to such behavior or its remnant.

We present snapshots of network dynamics induced by IL2 (IL2DL/IL2DR, IL2L/IL2R, IL2VL/IL2VR) neurons stimulation in the bottom panel of Fig. 4. Panel A. Notably, the network activates neurons that are located mostly on right side of the graph, which is a different pattern than forward and backward patterns. Most responsive motor neurons for such stimulation are RMG, RMH and RMED along with moderate responses within SMD and RMEL/RMER motor neurons. The oscillatory periods for these neurons are uniform around ~5.7s with different phases. Particularly, RMHL and RMHR neurons each produce oscillations nearly anti-phase to each other. For RMG neurons, the oscillation wave of RMGL always preceded that of RMGR, suggesting the phase displacement between oscillations of these two neurons. However, oscillations among 4 SMD motor neurons (SMDDL, SMDDR, SMDVL and SMDVR) as well as of three RME motor neurons (RMEL, RMER, RMED) were observed to be approximately in phase.

In the literature, these motor neurons are known to be involved with control of head muscles. RMG and RMH motor neurons innervate lateral four rows of head muscles while RME neurons innervate all eight rows of head muscles [3]. SMD motor neurons are also known to innervate head muscles involved with locomotion related to search behaviors such as omega-shaped turns under absence of food in the environment [40]. Remarkably, the visualization shows no response among the motor neurons associated with forward/backward locomotion (such as Ventricular Dorsal A, B and D) but only activated motor neurons modulating head muscles. Such results suggest that the activation of IL2 neurons lead to periodic head movements with absence of locomotory behavior in the rest of the body. While this does not necessarily imply that such motor neuron pattern is linked to nictation, these observations provide particular hypotheses and insights about the relatively unknown sub circuit to be further studied empirically.

**Fig 5.**
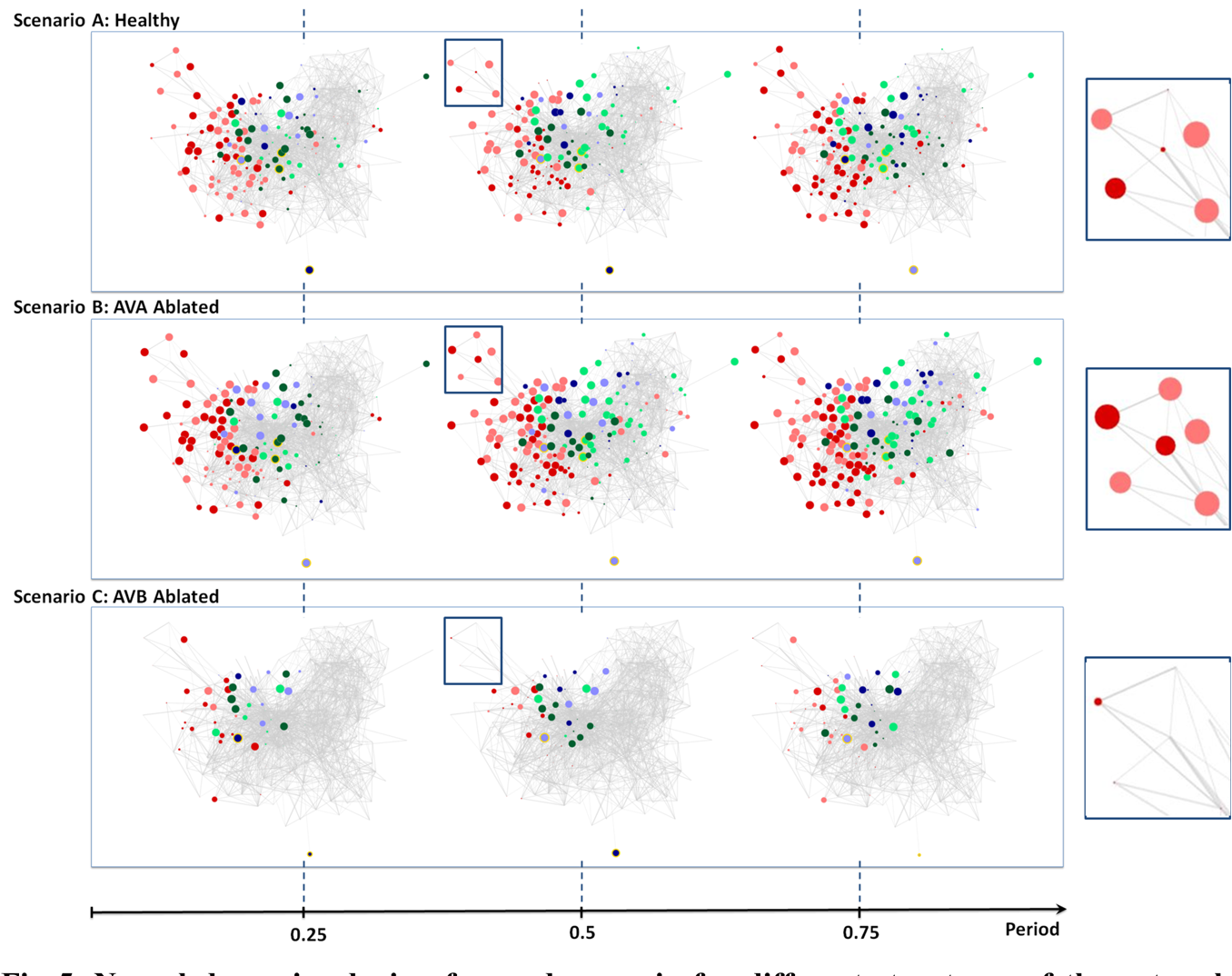
Neural dynamics during forward scenario for different structures of the network (non-ablated and ablated neurons). A: Three snapshots of dynamics with healthy network (i.e. no ablated neuron). B: Snapshots of dynamics when AVA inter neurons are ablated. C: Snapshots of dynamics when AVB inter neurons are ablated. All snapshots are taken at 25%, 50% and 75% into each dynamics’ oscillation period.

#### Scenarios: Ablation

For the validation of Neural Interactome’s application for investigation of network structural changes, we perform two ablations in conjunction with previously performed scenarios. In particular, we remove AVB and AVA interneurons from the network and repeat the posterior touch response scenario to observe their effects on the dynamics.

##### AVB ablation

According to the literature, the removal of AVB neurons impedes forward locomotion [25]. Indeed, we are able to confirm these experimental findings using Neural Interactome. Scenario C in Fig. 5 shows the three snapshots of full periodic cycle upon repeating the posterior touch response scenario with AVB neurons ablated. We observe that neural patterns involve far less neurons than that of healthy network (Fig. 5, Scenario A). In particular, we do not observe, through snapshots of full network as well as the local group of motor neurons, strong motor neurons responses in comparison to healthy structure.

The visualization does capture some weak oscillations within a small group of motor neurons; particularly in Ventricular Dorsal B (VB, DB). Oscillation amplitudes are far less than the healthy dynamics, however, they remain to be relatively in phase and maintain an oscillatory period of ~1.9 seconds. We are unable to find any oscillatory activity within Ventricular Dorsal type D (VD, DD) neurons, which were out of phase to the activity of (VB, DB) neurons in the healthy case. Acknowledging that the two modes oscillatory property is necessary for the worm to perform forward crawling motion [36], such observation confirms the experimental findings that the ablation of AVB neurons hinders the worm’s ability to perform forward motion.

##### AVA Ablation

Unlike the removal of AVB interneurons, experiments showed that the removal of AVA interneurons does not have impact on forward motion [25]. Scenario B in Fig. 5 shows snapshots of posterior touch response scenario with AVA neurons ablated. It is interesting to observe that the dynamics have slightly longer oscillatory period of about ~2.6 seconds. However, aside from that, the visualization shows that almost identical set of neurons are active as in the healthy scenario (compare with Fig 5, Scenario A). We are also able to confirm, using the review mode, that the dynamics continue to exhibit strong oscillations in (VB, DB) & (VD, DD) motor neurons, with (VD, DD) neurons being out of phase to (VB, DB) neurons. Thus our results for AVA ablation are consistent with experimental data in the literature.

Taken together our results show that Neural Interactome’s network visualization assists in confirming previously reported empirical results from the literature in effective manner, and further provides insights regarding structure and activity associated with examined responses.

## Discussion

In this paper, we present a new visual interactive method, which we call Neural Interactome for studying the dynamics and the structure of a neuronal system (*dynome*). While it is important to simulate the full *dynome* to study network functionalities, multiple simulations of the *dynome* are formidable, due to complexity in number of neurons, time and variations of stimulations. Here we demonstrate that the Neural Interactome approaches the problem through interactive real time interface to the *dynome* and therefore significantly simplifies these studies. In particular, we show the simplicity of stimulating various groups of neurons in the framework for testing experimental hypotheses, subsequently validate the *dynome* model, and obtain new hypotheses.

To elucidate the overall structure and functionalities of the framework, we first define key components: Interactive interface and backend neural integration. Next, we apply it to *Caenorhabditis elegans (C. elegans)* nematode, which connectome is resolved and the computational model describing both biophysical processes and interactions between neurons has been recently developed. We show that the framework provides novel possibilities to explore the worm’s network structure and its unique neural patterns subject to stimuli. In particular, we demonstrate the Neural Interactome’s capabilities using stimulations associated with touch response: stimulation of PLM/AVB neurons for posterior touch and ALM/AVA/AVD/AVE neurons for anterior touch, and stimulation associated with nictation behavior: stimulation of IL2 sensory neurons. In all three scenarios, we observe clear visual characteristics of the induced neural patterns. For example, using the review features, we are able to identify most responsive neurons and additional properties of dynamics such as oscillation period and phase on individual and population level. By comparing such observations with behavioral and neural descriptions in the literature, we demonstrate that our results are consistent with the empirical observations in regards of worm’s locomotion and suggest additional novel insights to be explored for relatively unknown sub circuit.

In addition, we demonstrate the effectiveness and usability of the neural ablation feature in the framework by ablating hub interneurons (AVA or AVB) in the network. AVB ablation leads to visualization with diminished activity in motor neurons associated with forward motion, as well as absence of characteristic out of phase oscillatory property required for such motion. The ablation of AVA interneurons, however, shows almost identical set of participating neurons as of the healthy network. We therefore believe that the framework has a potential to reveal other interesting functionalities through multiple ablation scenarios, and provide further insights about the role of the ablated neurons [42]. In experiments, preparation and execution of ablation consumes a lot of time and usually requires special equipment, e.g. optogenetics. On the contrary, Neural Interactome can produce initial analyses for numerous ablation scenarios within seconds. We thereby expect that the framework could be used as a pre experiment tool for mapping the scenarios to be explored empirically.

While *C.elegans* has well described connectome and founded models for dynamic neural data, it is still not a complete *dynome* of the worm’s nervous system with all possible biophysical neural interactions and connectivity at different developmental stages. We therefore designed the Neural Interactome to permit updates to both connectivity and dynamic models within the framework as they are further being refined in the future. Connectivity updates will require only a change in the connectivity matrices. Replacement of current model with more detailed one would require only the replacement of the model itself, while the synchronization method between the neural integration and interface will ensure that the computed values will be visualized properly. With such flexibility we expect that the framework will be similarly applicable to other neuronal systems: ranging from actual biological networks (such as that of *Drosophilla* medulla, the mouse retina, the mouse primary visual cortex) to artificial dynamic neural networks (e.g. Recurrent Neural Networks) and genetic networks [43]. We also plan to keep adding more features to the framework, to provide additional interaction possibilities with more detailed properties of the network, such as modification of individual synaptic or gap connections between a pair of neurons.

## Materials and Methods

In this section, we describe the materials and methods used for development of Neural Interactome and its application to *C.elegans* nervous system.

### Development Environment & Tools

We used two different programming languages for the development of Neural Interactome. In particular, we used Python for development of backend neural integration, and Javascript for frontend interactive interface. For establishing communication protocols between the interface and backend, we used flask-socketIO on python side and Socket.IO on javascript side. Both flask-socketIO and Socket.IO are libraries that allow for real time bi-directional communication between the client (frontend) and server (backend) through WebSocket protocols. In the context of Neural Interactome, they were used to establish the robust command ↔ data transactions between the interactive interface and backend neural integration.

Several third party libraries were used for each language as well. For Python, NumPy was extensively used for mathematical computations and manipulations of matrices. Several functions from SciPy were used to construct ordinary differential equation solver and solve the system of linear equations in regards of computation of neural quantities such as V-threshold values.

For Javascript, D3.js (Data-driven documents) platform was used to construct force-directed graph representation of neuronal network. For the main webpage development framework, we used AngularJS as it provides optimal functionalities for building dynamic, single page web apps (SPAs).

### V*_threshold_* Computation

At voltage equilibrium of the network, 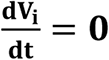 (Eqn. 2 for *C.elegans*). For computing network voltage values that satisfy such condition, we solve a following linear equation:

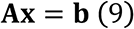

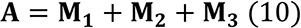

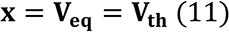

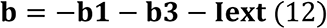

**M_1_** is a matrix of size N×N where N is the number of neurons (279 for *C.elegans*) with its diagonal terms populated with **−G^c^** (cell membrane capacitance).

**M_2_** is a diagonal matrix where diagonal term in i_th_row corresponds to 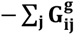.

**M_3_** diagonal matrix where its diagonal term in i_th_term corresponds to 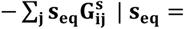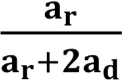.

**b_1_ = G^c^ * E_c_** where **E_c_** is a 1D vector of size N×1 in which all of its elements are ***E_c_***.

**b_3_ = G^s^ · (s_eq_ * E_j_)** where **E_j_** is a 1D vector of size N×1 that enlists the directionality of each neuron (0 if excitatory or −48mV if inhibitory).

**I_ext_** is the input stimuli vector where its i_th_ element determines the input current amplitude for i_th_ neuron.

### Parameters

The parameter values we used for our application of Neural Interactome to C.elegans nervous system are listed.

#### Dynome Visualization

For the maximum radius of the nodes in Eqn. 1, we used **R_max_** = 15.

#### Simulation Timescale

The following values were used for the temporal resolution of simulation and dynamic timescales for visualization.

**Temporal resolution:** 10ms

**Visualization rate (normal):** 100ms/s

**Visualization rate (during transition or ablation):** 40ms/s.

### Parameters for Neural Integration

The precise values of parameters for each connection described in Eqn. 2 ~ 6 are unknown. However, we assume reasonable values that are previously considered in the *C.elegans* literature [4], [11]. We assume each individual gap and synaptic junction has approximately equal conductance of **g** = 100pS [4], cell membrane conductance **G^c^** = 10pS, and membrane capacitance **C** = 1.5pF [4]. We take leakage potential **E_cell_** = −35mV while reversal potential **E_j_** = 0mV for excitatory synapses and −48mV for inhibitory synapses [11]. For the synaptic activity variable, we take 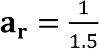, 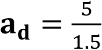 and width of the sigmoid **β** = 0.125mV^−1^ [11]. Also for the initial condition of the membrane voltages V and synaptic activity variable s, we sample the normal distribution of µ = 0 and σ = 0.94 with size 279 * 2 (for both V and s) and multiply by 10^−4^.

### Parameters for Synchronization

The optimal values for **Δt, t_buffer_**, and internal refractory period τ in Eqn. 7 depend on computing power. However, we found the parameters **Δt = 50ms, t_buffer_ = 100ms** and **τ = 50ms** (in actual time) to be of reasonable default values which achieve both computational efficiency and synchronization between the interface and backend. Note that **Δt** and **t_buffer_** are in simulation timescale while τ is measured in computer’s internal timer.

### Parameters for Stimuli transition

We use ***t_offset_*** = 150*ms*, and **r** = 0.025 for Eqn. 8. We have found these values to be the optimal choices as its transition curve isn’t too aggressive to induce abrupt shift in *dynome* dynamics, but also reasonably short enough that the visualization rate doesn’t stay in slow mode for too long. Given the value of **r**, the time it takes for complete transition from one stimulus amplitude to the other is approximately **t_buffer_** * 2. Thus for our choice of parameters for *C. elegans* simulations, the transitional period is around 300ms.

## Footnotes

1 Web interface available at http://neuralcode.amath.washington.edu/neuralinteractome

